# Life history trade-offs in ectotherms impart a pseudo-hormetic response, not hormesis

**DOI:** 10.1101/337279

**Authors:** Malcolm L. McCallum, Michelle Mary, Stanley E. Trauth

## Abstract

All organisms are faced with survival and fitness challenges. However, differences in how ectotherms and endotherms deal with these challenges causes confusion when theoretical explanations are proposed. Herein, an immunochallenged, immature ectothermic vertebrate increased growth in the face of immunochallenge, reminiscent of up-regulated physiological efficiency termed hormesis. The immunochallenged subjects increased food intake relative to control animals, a largely ignored possibility in previous studies. This likely led to an energy surplus that fueled additional growth. Although there was increased resource demand from the immune response that exceeded internal stores, the acceptably sized food items contained more resources than immunologically-driven demand required. We theorize that because ectotherms lack significant internally-stored resources compared to endotherms, they must feed to fuel increased physiological demand. This can lead to excess resource intake because the minimum acceptably sized prey contains more available resources than upregulation required. This creates a pseudo-hormetic hormetic response, fueled by excess food intake rather than significant improved physiological efficiency. Further, we speculate that lifespan and/or maturity may interact with resource management ectotherms, though our data are inconclusive on this matter. Ultimately, our data suggest additional growth when ectotherms face stressors is a pseudo-hormetic response stemming from increased food intake instead of upregulated physiological efficiency.

## Introduction

Life history components are the outcomes from exposure to environmental stressors over evolutionary time. The Law of the Minimum states these components are regulated by the availability of one or a few factors in short supply; whereas, resources in excess may go unused (Liebig 1840). The Principle of Allocation (Levins 1968) suggests an organism’s resource budget is distributed among various life history components loosely categorized among maintenance, growth, reproduction, activity, social behavior and immunity (McNab 2002; Stearns 1992; Ricklefs and Wikelski 2002):

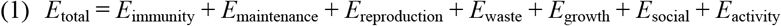

When challenged, resources are shifted among components in accordance with these two rules, based on efficiency and need (Parrish and Edelstein-Keshet 1999; Sheldon and Verhulst 1996; Lochmiller and Deerenberg 2000). Among the more formidable challenges an organism faces is its battle with pathogens, parasites and environmental stressors (Fair and Ricklefs 2002; Schmid Hempel and Schmid Hempel 2011; Hawley and Altizer 2011). These adversaries cause stress by physically and chemically interfering with normal biological functions and disrupting homeostasis (Chrousos and Gold 1992). Some of the many strategies for dealing with these opponents include non-specific immunity (Anderson 1993; Bosch et al. 2003; Fair and Ricklefs 2002), specific immunity (Fearon et al. 1996), shedding diseased or injured body parts (Scott and Jakob 1999; Randall 1981), or in many short-lived species, simply coping with them (Kapahi et al. 1999). As defenses are activated, resources must be drawn from other areas of life history (Sheldon and Verhulst 1996; Norris and Evans 2000; Schulenburg et al. 2009) or their availability increased (Le et al. 1998; Zulkifli et al. 2000; Zellner et al. 2006). In some cases, organisms faced with immunochallenge have drawn resources from other areas of life history to implement defense (Lochmiller and Deerenberg 2000; McCallum and Trauth 2007; Sheldon and Verhulst 1996). In many cases; however, low-dose immunochallenge improves growth (e.g. hormesis) and extends lifespan without clear trade-off effects (Alexander 1974; Reed et al. 1986). Life history strategies are critical determinants of how organisms deal with stress (Stark et al. 2004).

The theory of hormesis suggests stressors at low and often intermittent exposures may stimulate growth, despite inhibiting growth at higher doses (Calabrase and Baldwin 1998; Mattson 2008). Hormesis was hypothesized in the 1800s (Schulz 1888); however, its origins extend to the 16^th^ Century (Stebbing 1982; Calabrase and Baldwin 1998). Since then, thousands of studies have revealed hormetic responses (Calabrese and Baldwin 1998) to ischemic preconditioning (Arumugam et al. 2006), exercise (Radak et al. 2005), dietary energy restriction (Qui et al. 2010), exposure to low doses of chemicals (McCallum et al. 2013; Li and Chen 2005) or pathogens (McClure et al. 2014), and a multitude of other stressors (Calabrese et al. 2007; Gems and Partridge 2008; Schreck 2010). Even many vitamins follow a hormetic response in that they are beneficial in low doses, but detrimental in excess (Mattson 2008 c). Hormesis is particularly relevant to environmental toxicology, biomedical and clinical sciences, and immunology (Calabrase and Baldwin 2003; Mattson 2008 c).

Hormesis can be at the cellular and/or the organismic level, leading to confusion for investigators (Calabrase and Baldwin 2002). The molecular mechanisms mediating hormesis involve an array of enzymes and transcription factors that increase production of cytoprotective and restorative proteins including growth factors, phase 2 and antioxidant enzymes, and protein chaperones (Mattson 2008). The ultimate cause of growth lies in either a biopositive effect of or overcompensation for stress (Calabrese and Baldwin 2002). The Direct stimulation hypothesis proposes a vastly regulated system with major resource constraints that reflect normal, modulatory physiological dynamics (Calabrase and Baldwin 2002). The overcompensation hypothesis functionally links hormesis to homeostasis, suggesting a “highly regulated, optimization process providing additional adaptive equity…as a biological insurance policy” (Calabrase and Baldwin 2002; Calabrase 2008). Resolution of this issue remains elusive (Stebbing 1982; Calabrase and Baldwin 1998; 2002). Regardless, hormesis is an adaptive, dose-specific response induced by at least one of these options (Calabrase and Baldwin 2002). Further, it is a fundamental concept of evolutionary theory (Mattson 2008) that challenges the most basic precept in toxicology, the dose-response relationship, and its two traditional paradigms, the threshold and non-threshold models (Calabrese 2005; Mattson 2008).

The ecology of the pathogen and chemical induced stress response in amphibians is of particular interest because of the prominent role immunity (Carey et al. 1999); *Batrachochytrium dendrobatidis* (Fisher et al. 2009; Ramsey et al. 2010), ranavirus (Greer et al. 2005; Daszak et al. 1999; Schloegel et al. 2009) and various chemical contaminants (Blaustein et al. 2002; Hayes et al. 2006; Jones et al. 2010) have played in their global-scale decline (Houlahan et al. 2000; Collins and Storfer 2003; McCallum 2007; McCallum 2015). The amphibian immune system is similar to that of fishes and less sophisticated than that found in mammals and birds (Pastorett et al. 1998; Manning and Turner 1976). Like these other taxa, amphibians have a well-developed non-specific immunity composed of antioxidants, chemical antagonists, complement, interleukins, and physical barriers (Pastorett et al. 1998; Manning and Turner 1976). Cell mediated response is well-developed (Manning and Turner 1976). Amphibian specific immunity is less elaborate than in higher taxa, with the humoral response limited to a primary antibody class, Immunoglobulin M (IgM) and a secondary class, Immunoglobulin Y (IgY) (Manning and Turner 1976). The former is found in all vertebrate and most invertebrate taxa, the latter is unique to amphibians, and similar in structure and function to Immunoglobulin G (IgG) found in mammals. Unlike birds and mammals, the kinetics of amphibian physiology, and therefore of its immune response, are keyed to the ambient temperature (Ribas et al. 2009; McCallum 2003; Manning and Turner 1976). In cold winters, amphibian immunity appears limited to standing levels of non-specific defenses and titers that were produced and stored during warmer weather.

As the lowest terrestrial vertebrate class, the Amphibia constitute a major evolutionary step in the development of stress responses related to the primordial invasion of the land. Like other organisms, Amphibia express physiological trade-offs when faced with challenges to different life history components, including immunity (McCallum and Trauth 2007; Garner et al. 2009; Gervasi and Foufopoulos 2008). However, it is not clear how food consumption interacts with immunity and possible trade-offs, while amphibians deal with stressors.

Hormetic responses to stressors by amphibians appear in some studies (Wijesinghe 2012; James and Little 2003; Perez—Coll et al. 2008) but not others (Miyachi et al. 2002; Urine et al. 2004; Endler et al. 2015). Considering the known interactions between food intake, growth, and stress/immune responses in other organisms (especially homeotherms), we asked if immunochallenge of an immature ectherm, the Green Tree Frog *(Hyla cinerea*), might also draw resources from growth to fuel immunity. We predicted that if this happened, those provided a novel antigen should grow less during the study than others injected with a non-antigenic control.

## Materials and Methods

Wild-caught Green Tree Frogs (n = 50) were assigned to an experimental and control group. Frogs were housed individually in 4-L plastic containers at 22 °C and with a 12 hr photoperiod. Each housing unit included a 5cm cube of natural sponge saturated with water to maintain humidity. Each frog was weighed at the beginning and end of the study. Each frog was fed a single juvenile House Cricket (*Acheta domesticus*) once a week. The cricket was weighed and placed in with the frog. If the cricket remained in the bag after one-week it was removed and tallied as uneaten, then replaced with a new cricket. Frogs were inspected daily, and housing was cleaned at least weekly.

We elicited immune responses in the experimental group with washed whole sheep blood as a novel antigen. This is known to produce detectable IgM responses in amphibians (Manning and Turner 1976; McCallum 2003; McCallum and Trauth 2007). Use of a novel antigen avoids toxin accumulation associated with pathogens and secondary immune responses related to previous exposure (Garvey et al. 1977). Whole sheep blood was rinsed three times in 0.85% saline. Then, it was volumetrically diluted to 100 ml with 0.85% saline to form a 1% solution of sheep blood cells (approximately 2 × 10^8^ cells/ml) according to methods outlined in Campell et al. (1970). Light scatter of the blood solution was read on a Spec 20 GenesysTM (Spectronic Instruments) at 560 nm using 0.85% saline as the blank control. The absorbance (actually light scatter) for the blood solution (2.148) was used to calibrate the concentration of cells prior to each weekly treatment.

Experimental frogs were injected weekly with 0.1 ml of blood solution and control frogs were injected with 0.1 ml of 0.85% saline. Frogs received injections into their dorsal lymph sac. This regiment continued weekly for three weeks, and the study was ended at the end of day 21. Mortality, growth, and food intake were compared between treatments and to the earlier results with the short-lived Blanchard’s Cricket Frog, *Acris blanchardi* (McCallum and Trauth 2007). Green Tree Frogs may live > 5 yrs and breed several times (Encyclopedia of Life. Available from http://www.eol.org. Accessed 3 May 2017); whereas, cricket frogs typically live < 2 yrs and seldom survive past participation in their first breeding season (McCallum et al. 2011). We used α < 0.05 as significant, α > 0.1 as not significant, and 0.5 < α < 0.1 marginally significant when applying decision theory.

## Results

Survivorship did not differ between treatments (***X***^2^ = 0.50, df = 1, P = 0.4795). The experimental group grew more than the control group (U = 201, P = 0.048). The control group’s weight remained unchanged through the study (T = −0.33, P = 0.743). Average food intake by the experimental group was significantly more than the control group (U = 81, P = 0.001). One frog in the control group refused all food in the study and it lost 0.01 gLW.

The experimental group consumed 19.5% more of their crickets than did the control group. Experimental animals consumed ~36% more energy during the study. Frog tissue is 81.7% water (18.3% DM) and contains 1.638 kcal/gDM (Cummins and Wyucheck 1971). Although this value excludes bony elements, we used this reasonable guideline to show immunochallenged frogs used roughly 0.046 kcal for growth; whereas, the control group had no net gain or loss in bodyweight. Therefore, estimating *E*_total_ based on the energy content of the mass of crickets consumed, we removed *E*_growth_ from *E*_total_ of the experimental group:

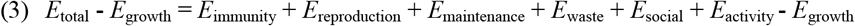

Then, assuming *E*_maintenance + social + reproduction + activity + waste_ was similar between the experimental and control groups, we could remove it from the equation and calculate an estimate for *E*_immunity_ for the experimental group:

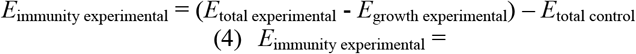

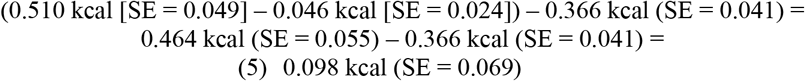

The cost of immunity while ignoring minor differences in waste production associated with additional food processing/digestion by the experimental group was roughly 0.098 kcal during the life of this study, the excess energy from food was shunted to additional growth.

**Table 1.**
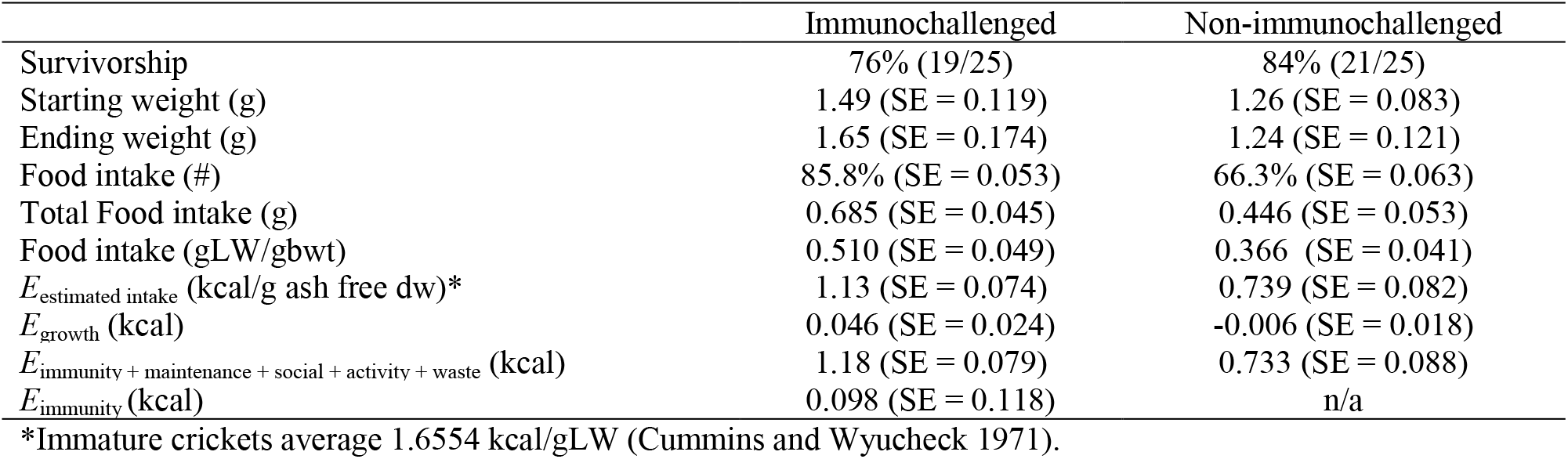
Responses of Green Tree Frogs (*Hyla cinerea*) to immunochallenge.

## Discussion

Green Tree Frogs responded to immunochallenge by increasing their energy intake, compared to members of the control group, to fuel their immune response. Because their energy deficit was sufficient to drive hunger, but the available prey contained more energy than needed, experimental frogs gained weight.

### Energy reserves and growth

Trade-off theory suggests that resources are shifted among life history components based on need when resources are limited (Gadgil and Basert 1970; Charnov and Krebs 1973; Stearns 1992). Our study shows that when internal resource budgets are exceeded, ectotherms may increase intake to supplement growing needs (MacKay 1985; Kooka et al. 2009). Further, this supplementation may result in a resource surplus that the body can redistribute to growth or potentially other life history components. Consequently, physiological trade-offs might be driven by internal resource banks, but ectotherms have mechanisms available to supplement internal resources when demand grows (MacKay 1985). Hence, this reality allows modification of the resource budget represented by equation (1) to encompass this adaptive consequence for life history:

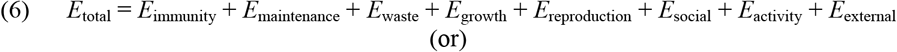

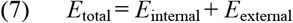

Where *E*_internal_ = the resources stored in the body, and *E*_external_ = the increased resources an organism delivers via raised feeding levels. *E*_external_ should be limited by the kinds and number of foods available (Holling 1959; Gross, et al., 1993; Spalinger and Hobbs 1992), and the organism’s capacity to utilize those food items:

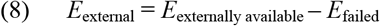

Where *E*_externally available_ represents all potential prey or food items in the vicinity and *E*_failed_ is all foods that are not used, regardless of reason (e.g., lost opportunity, anti-predator defenses, predator familiarity, inability to digest food components, etc.).

Food intake, endocrine function and immune response in mammals involve leptin. Leptin is produced by adipose tissue and mutations of the genes coding for it and its receptors can lead to obesity (Zhang et al. 1994; Coleman 1978; Tartaglia et al. 1995). Some believe leptin signals the nutritional status to regulate homeostasis of energy balance and reduce food intake in mammals (Faggioni et al. 2001). Leptin is regulated during the acute-phase response, is acutely increased by inflammatory and infectious stimuli and is related to pathophysiological changes such as hypoglycemia and anorexia in humans (Gabay and Kushner 1999). Amphibians, like mammals, express a single ortholog of leptin (Boswell et al. 2006; Crespi and Denver 2006; Londraville et al. 2014); however, amphibians store far less fat than do mammals (Brown 1964). Like mammalian leptin, the amphibian form can function as an anorexigen, but its function in vertebrates varies widely (Faggioni et al. 2001). In fact, studies on Amphibian lectin are few. Woodchuck (*Marmota marmot*) Leptin titer and body mass are out of phase during hibernation (Concannon et al. 2001). In humans, there is enormous variability in body mass index and leptin titer (Cnop et al. 2002). Most fishes increase leptin or leptin mRNA during fasting periods, then levels subside after feeding (Fuentes et al. 2013). A mutant leptin gene in Medaka increases appetite and growth rate but does not increase fat stores (Chisada et al. 2014). Leptin may be directly linked with the hypothalamo-pituitary-adrenal/interrenal axes (Roubos et al. 2012) and their interactions with food intake (Ahima et al. 2000), the stress response (Glasow and Bornstein 2000; Heiman et al. 1997), and immunity (Faggioni et al. 2001). Because immunochallenged Green Tree Frogs grew more rapidly than the control group, and that growth was fueled by increased food intake like seen in fishes; one can speculate that leptin may play a similar, unique, and little studied role in physiological trade-offs for this species, if not ectotherms in general.

### Implications for Hormesis

Resolution of the ultimate cause of hormesis has been difficult and current dogma suggests involvement of a biopositive effect of or overcompensation for stress (Stebbing 1982; Calabrese and Baldwin 1998; 2002). However, our results provide a tangent possibility. In some cases, pseudo-hormesis may stem simply from increased food intake when total metabolic costs grow due to stress on one or more body systems. Clearly, if an animal increases feeding behavior when faced with minor stress, this violates the overcompensation hypothesis because even though major resource constraints exist, these will only persist if food is unavailable.

This explanation does not seem sufficient for many mammals, including humans, where lectin suppresses appetite and therefore food intake during the stress/immune response (Faggioni et al. 2001). Endotherms maintain very high energy budgets, have lower production efficiencies, and store many resources internally to maintain homeostasis (Hamilton 1973; Young 1976; Heinrich 1977); whereas, poikilotherms are thermal conformers with higher production efficiencies, thus requiring fewer stored resources to survive (Huey and Slatkin 1976; Lavigne 1982; Pianka 1985; 1986; 1993). Storage of excess energy is less critical for poikilotherms under normal circumstances, but more risky when faced with resource-consuming stress (Lavingne 1982; Sheridan 1994). Ectotherms that invest in life histories (e.g. parental care, migration) tend to store more resources than those that do not (Cho et al. 1982; Sheridan 1994; Arrington et al. 2002; Stallings et al. 2010). This difference in stored resource banks should cause excessively-stressed ectotherms to exhaust internal resources quicker than would similarly stressed endotherms. Endotherms have much higher resource demands, but store more excess energy (Young 1976: Pond 2011). This may allow them to shift a larger bulk of resources, due to internal availability, from one system to another than ectotherms can when facing crises (Young 1976). Once comparatively meager resources are exhausted by ectotherms, they must either increase intake or succumb to stress (Wieser 1991; Griffiths and Kirkwood 1995; Lazzaro and Galac 2006). Intake is limited by the relative size of a food item and mouth size (Toft 1980; Brodie and Formanowicz 1983; Dabrowski and Bardega 1984; Forsman 1996). When physiological stress is low to moderate, ectotherms feed upon the appropriate-sized food whether it has insufficient, sufficient, or excess resources in its bank. Once the immediate needs of the stressed life history component are satisfied, any excess resources can be shifted to other systems for use. *Hyla cinerea* fed on crickets that contained more resources than they required for the increased immune response, so they shifted those excess resources to growth; whereas, an endotherm may have stored those in adipose tissue. Similar findings were reported with Largemouth Bass (*Micropertus salmoides*) where these poikiliotherms shunted excess resources to growth and reproduction rather than storage (Garvey and Marschall 2003). Conversely, Atlantic Silversides (*Menidia menidia*) do not demonstrate trade-offs with growth, instead shifting resources with burst speed (Billerbeck et al. 2000). However, otherwise unstressed endotherms faced with food deficits exhaust resources rapidly when food is not regularly available (Stanier et al. 1984).

Hormesis in amphibian tadpoles is said to arise from stress sourced to predator exposure that led to increased cortisol production (Maher et al. 2013). However, tadpoles feed on detritus and particulates in water as well as bacterial film and algae (McDiarmid and Altig 1999). Tadpoles could up-regulate feeding behavior just by filter feeding, which could easily go unobserved. In fact, why well-fed mother *Daphnia* sp. that were smaller grew and reproduced better than food-starved mothers went unexplained until it was revealed that offspring of food-starved females of *Daphnia magna* spent less time feeding. Food intake led to differences in growth and stemmed from lower investment of resources in the offspring by these ectotherms (Garbutt and Little 2014). Hormesis was reported in bacteria (Christofi et al. 2002), *Caenorhabditis elegans* (Cypser and Johnson 2002), Drosophila melanogaster, *Plutella xylostella* (Ebrahimi et al. 2015), *Daphnia* sp. (Zalizniak and Nugegoda 2006; Li and Tan 2011; Flaherty and Dodson 2005), juvenile mussels (Pandolfo et al. 2010) and an assortment of other small ectotherms. All suffer from the same issue: Did growth arise from new efficiencies as suggested in endotherms, or simply from eating more food as seen in *D. magna*? Directly observing changes in food intake with these animals is difficult at best. Hence, if this ectothermic effect on food intake when facing minor stress took place, leading to a pseudo-hormetic response, investigators might not ever notice. We observed a similar response in Green Tree Frogs, supporting the notion that hormetic responses in ectotherms do not follow the same pattern as they do in endotherms.

Based on the traditional definition of hormesis, Green Tree Frogs do not respond hormetically to a stressor. Rather, it appears they supplement body resources via additional feeding to fuel responses. The improved body growth (hormesis) is simply a side-effect of excessive nutrient intake due to the ratio between size appropriate prey and the energy deficit stressors are causing. It is likely that many ectotherms deal with these trade-offs in the same manner, but this behavior is masked by food particle size and the body size of the subject organism.

### Trade-offs between reproduction and body size

The cost of reproduction remains among the most prominent life history trade-offs studied and involves two well-researched components, costs paid in survival and in future reproduction (Stearns 1989; Williams 1957, 1966; Hirshfield and Tinkle 1974; Calow 1979). Whether an organism arrives at a larger body size via hormesis or some other path, this provides fitness advantages over conspecifics that are smaller. First, there is a higher probability of survival to first reproduction for larger individuals than smaller ones (Brodie and Formanowicz 1983; Anderson 1988; Lindstrom 1999). Further, larger females can produce larger clutches of eggs (Darwin 1874; Shine 1988) and sometimes larger eggs (Shine 1988; Congdon and Gibbons 1987), both which have distinct reproductive advantages. Larger males are more competitive where intrasexual competition exists, possibly even entering the breeding population earlier than is typical for their ages (Meneses 2007; Jones and Hutchings 2001; Arak 1983). Early emergent male, but not female Blanchard’s Cricket Frogs (*Acris blanchardi*) can reach physiologic maturity the same summer they hatch; however, social factors likely force this cadre of offspring to wait until the following breeding cycle to participate in the chorus (McCallum et al. 2011). In fact, smaller and younger frogs often participate as satellite males surrounding a more competitive, large, older male, functionally parasitizing him for mates (Wells 1977; McCallum et al. 2006; Leary et al. 2004), although direct fighting can also take place between males (Wagner 1989). This suggests that there may be a selective advantage for ectotherms that are exposed to stressors early in life, assuming the stress response leads to a sufficient resource drain to elicit hunger, but a minimum prey size that provides resources in excess of that deficit. Such conditions would be highly adaptive to ectotherms that chose to leave offspring under conditions that were suitable, but not completely ideal for development. Clearly, the Green Tree Frogs that experience a certain low-level threshold of stress will grow faster, and be more competitive in the breeding chorus, leading to greater fitness than those exposed to no stress or high levels of stress simply because they eat more food when under a trade-off induced energy deficit.

### Life span or maturity may play a role in life history trade-offs between immunity and reproduction

Blanchard’s Cricket Frog is an annual species in this portion of its range (McCallum et al. 2011); whereas, the Green Tree Frog lives several years and participates in multiple breeding choruses during its lifetime. This difference in species-specific lifespan may explain differences in how trade-offs with immunity are handled and warrant further investigation.

When faced with immunochallenge, an organism may shift resources between survival and reproduction to maximize fitness (Stearns 1989). Although both Green Tree Frogs in this study and Blanchard’s Cricket Frogs from the previous study (McCallum and Trauth 2007) were both about the same age (6 – 10 mo old), they were not at the same reproductive maturity level. Green Tree Frogs were in the growth phase in preparation for life-time reproduction; whereas, Blanchard’s Cricket Frogs were in the midst of peak reproductive activity of their only breeding season. Green Tree Frogs opted to invest excess resources in growth, which improved fitness by enhancing body size, hence competitiveness in the breeding chorus and capacity to hold eggs. Immunochallenged Blanchard’s Cricket Frogs did not invest excess resources into growth, if any surplus was present. Instead, it appeared they either shunted the excess resources into immunity or some other life history component. They did not appear to shift resources to reproduction because they did not retain pigments associated with reproduction throughout the series of immunochallenges, and tissue atrophy in the testes was observed (McCallum and Trauth 2007; McCallum 2003). These species may trade-off resources differently because each faces different reproductive opportunity. By shifting resources to growth, Green Tree Frogs maximize future fitness along their much lengthier lifespan. By exhausting resources now, Blanchard’s Cricket Frog maximizes current fitness because there will be no future opportunity for reproduction. These opposing life history choices lead to drastically different opportunities and risks related to reproduction, leading to the evolution of different physiological responses to immunochallenge.

### Trade-off theory, hormesis, and ectothermy

Although organisms evolve common strategies for dealing with stressors, key differences among life histories can result in divergent responses. For the endotherm, hormesis is certainly a response stemming from more efficient use of stored resources. However, ectotherms trade-off resources differently because of their vastly lower available internal storehouses evolved due to metabolically lower resource demands compared to endotherms. They rapidly exhaust internal resources and must shift to external sources. When that happens, the deficit driving hunger is exceeded by the gape size limitation placed on minimum acceptable prey size. They must intake more resources than they need. Without massive storehouses of adipose tissue, ectotherms shunt resources to other life history components to maximize fitness. In the case of longer lived immature ectotherms, a shift to growth is more adaptive. For short-lived reproductive ectotherms, it’s now or never. Growth advantages realized after the breeding chorus are useless. Maximum fitness is realized by reproducing now. Therefore, we see disparate results in how long-lived Green Tree Frogs and short-lived Blanchard’s Cricket Frog respond to immunochallenge and shunt resources.

## Literature Cited

Allen ME (1989) Nutritional aspects of insectivory. Doctoral Dissertation. Michigan State University, East Lansing, MI, U.S.A.

Alexander RD (1974) The evolution of social behavior. Annual Review of Ecology and Systematics 5:325 – 383.

Anderson RS (1993) Modulation of nonspecific immunity by environmental stressors. Chapter 17: Pathobiology of Marine and Estuarine Organisms. Pp. 483 – 510 In Couch JA, Fournie JW (eds.). Pathobiology of Marine and Estuarine Organisms. CRC Press. Boca Raton, Florida, U.S.A.

Anderson JT (1988) A review of size dependent survival during pre-recruit states of fishes in relation to recruitment. Journal of Northwest Atlantic Fishery Science 8:55 – 66.

Arak A (1983) Sexual selection by male-male competition in natterjack toad choruses. Nature 306:261 – 262.

Arrington DA, Winemiller KO, Loftus WF, Akin S (2002) How often do fishes “run on empty”? Ecology 83:2145 – 2151.

Beamish RJ, Mahnken C, Neville CM (2004) Evidence that reduced early marine growth is associated with lower marine survival of coho salmon. Transactions of the American Fisheries Society 133:26 – 33.

Billerbeck JM, Schultz ET, Conover DO (2000) Adaptive variation in energy acquisition and allocation among latitudinal populations of the Atlantic Silverside. Oecologia 122:210 – 219.

Blaustein AR, Kiesecker JM (2002) Complexity in conservation: Lessons from the global decline of amphibian populations. Ecology Letters 5:597 – 608.

Bosch JA, de Geus EJC, Veerman EJ, Hoogstraten J, Amerongen AVN (2003) Innate secretory immunity in response to laboratory stressors that evoke distinct patterns of cardiac autonomic activity. Psychosomatic Medicine 65:245 – 258.

Boswell T, Dunn IC, Wilson PW, Joseph N, Burt DW, Sharp PJ (2006) Identification of a non-mammalian leptin-like gene: Characterization and expression in the tiger salamander *(Ambystoma tigrinum*). General and Comparative Endocrinology 146:157 – 166.

Brodie Jr. ED, Formanowicz Jr. DR (1983) Prey size preference of predators: Differential vulnerability of larval anurans. Herpetologica 39:67 – 75.

Brown Jr. GB (1964) The metabolism of Amphibia. Pp. 1 – 98 in Moore JA, Physiology of the Amphibia. Academic Press. New York, NY, U.S.A.

Calow P (1979) The cost of reproduction – a physiological approach. Biological Reviews 54:23 – 40.

Charnov EL, Krebs JR (1973) On clutch sieze and fitness. Ibis 116:217 – 219.

Calabrese EJ (2005) Paradigm lost, paradigm found: The re-emergence of hormesis as a fundamental dose response model in the toxicological sciences. Environmental Pollution 138:378 – 411.

Calabrese EJ (2008) Overcompensation stimulation: A mechanism for hormetic effects. Critical Reviews in Toxicology 31:425 – 470.

Calabrese EJ, Baldwin LA (1998) Hormesis as a biological hypothesis. Environmental Health Perspectives 106 (Suppl 1):357 – 362.

Calabrese EJ, Baldwin LA (2002) Defining Hormesis. Human & Experimental Toxicology 21:91 – 97.

Calabrese EJ, Baldwin LA (2003) Toxicology rethinks its central belief. Nature 421:691 – 692.

Calabrese EJ, Bachmann KA, Bailer AJ, Bolger PM, Borak J, Cai L, Cedergreen N, Cherian MG, Chiueh CC, Clarkson TW, Cook RR (2007) Biological stress response terminology: integrating the concepts of adaptive response and preconditioning stress within a hormetic dose – response framework. Toxicology and Applied Pharmacology 222:122 – 128.

Carey C, Cohen N, Rollins-Smith L (1999) Amphibian declines: An immunological perspective. Developmental & Comparative Immunology 23:459 – 472.

Chisada S, Kurokawa T, Murashita K, Ronnestad I, Taniguchi Y, Toyoda A, Sakaki Y, Takeda S, Yoshiura Y (2014) Leptin receptor – deficient (knockout) Medaka, *Oryzias latipes*, show chronical uregulated levels of orexigenic neuropeptides, elevated food intake and state specific effects on growth and fat allocation. General and Comparative Endocrinology 195:9 – 20.

Cho CY, Slinger SJ, Bayley HS (1982) Bioenergetics of salmonid fishes: Energy intake, expenditure and productivity. Comparative Biochemistry and Physiology 73B:25 – 41.

Christofi N, Hoffmann C, Tosh L (2002) Hormesis responses of free and immobilized light-emitting bacteria. Ecotoxicology and Environmental Safety 52:227 – 231.

Chrousos GP, Gold PW (1992) The concepts of stress and stress system disorders: Overview of physical and behavioral homeostasis. Journal of the American Medical Association 267:1244 – 1252.

Coleman DL (1978) Obese and diabetes: Two mutant genes causing diabetes-obesity syndromes in mice. Diabetologia 14:141 – 148.

Collins JP, Storfer A (2003) Global amphibian declines: Sorting the hypotheses. Diversity and Distributions 9:89 – 98.

Concannon P, Levac K, Rawson R Tennant B, Bensadoun A (2001) Seasonal changes in serum leptin, food intake, and body weight in photoentrained woodchucks. American Journal of Physiology: Regulatory, Integrative, and Comparative Physiol. 281:R951 – 959.

Congdon JD, Gibbons JW (1987) Morphological constraint on egg size: A challenge to the optimal egg size theory? Proceedings of the National Academy of Sciences of the United States of America 84:4145 – 4147

Constantini D (2014) Oxidative stress and hormesis in evolutionary ecology and physiology: A marriage between mechanismtic and evolutionary approaches. Springer. https://link.springer.com/book/10.1007/978-3-642-54663-1.

Cnop M Landchild MJ, Vidal J Havel PJ, Knowles NG, Carr DR, Wang F, Hull RL, Boyko EJ, Retzlaff BM, Walden CE, Knopp RH, Kahn SE (2002) Diabetes 51:1005 – 1015.

Crespi EJ, Denver RJ (2006) Leptin (ob gene) of the South African Clawed Frog *Xenopus laevis*. Proceedings of the National Academy of Sciences 103:10092 – 10097.

Cummins KW, Wyucheck JC (1971) Caloric equivalents for investigations in ecological energetics. Mitteilungen Communications No. 18. Internationale Vereinigung Für Theoretische und Angewandte Limnologie. 158 pp.

Cypser JR, Johnson TE (2002) Multiple stressors in *Caenorhabditis elegans* induce stress hormesis and extended longevity. The Journals of Gerontology Series A: Biological Sciences and Medical Sciences 57:B109 – B114.

Dabrowski K, Bardega R (1984) Mouth size and predicted food size preferences of larvae of three cyprinid fish species. Aquaculture 40:41 – 46.

Darwin CR (1874) The edscent of man, and selection in relation to sex. 2^nd^ ed. Appleton, New York, NY, U.S.A.

Daszak P, Berger L, Cunningham AA, Hyatt AD, Green DE, Speare R (1999) Emerging infectious diseases and amphibian population declines. Emerging Infectious Diseases 5:735.

Donato AJ, Walker AE, Magerko KA, Bramwell RC, Black AD, Henson GD, Lawson LA, Seals DR (2013) Life-long caloric restriction reduces oxidative stress and preserves nitric oxide availability an function in arteries of old mice. Aging Cell 12:772 – 783.

Endler PC, Scherer-Pongratz W, Harrer B, Lingg G, Lothaller H (2015) Amphibians and ultra-high diluted thyroxine – further experiments and re-analysis of data. Homeopathy 104:250 – 256.

Fair JM, Ricklefs RE (2002) Physiological, growth, and immune responses of Japanese quail chicks to the multiple stressors of immunological challenge and lead shot. Archives of Environmental Contamination and Toxicology (42:77 – 87).

Fearon DT, Locksley RM (1996) The instructive role of innate immunity in the acquired immune response. Science 272:50.

Fisher MC, Garner TW, Walker SF (2009) Global emergence of *Batrachochytrium dendrobatidis* and amphibian Chytridiomycosis in space, time, and host. Annual Review of Microbiology 63:291 – 310.

Forsman A (1996) Body size and net energy gain in gape-limited predators: A model. Journal of Herpetology 30:307 – 319.

Fuentes EN, Safian D, Einarsdottir IE, Valdes JA, Elorza AA, Molina A, Bjornsson BT (2013) Nutritional status modulates plasma leptin, AMPK and TOR activation, and mitochondrial biogenesis: Implications for cell metabolism and growth in skeletal muscle of the fine flounder. General and Comparative Endocrinology 186:172 – 180.

Gadgil M, Bossert W (1970) Life history consequences of natural selection. American Naturalist 104:1 – 24.

Garner TWJ, Walker S, Bosch J, Leech S, Marcus Rowcliffe J, Cunningham AA, Fisher MC (2009) Life history tradeoffs influence mortality associated with amphibian pathogen *Batrachochytrium dendrobatidis*. Oikos 118:783 – 791.

Garvey JE, Marschall EA (2003) Understanding latitudinal trends in fish body size through models of optimal seasonal energy allocation. Canadian Journal of Fisheries and Aquatic Sciences 60:938 – 948.

Gems D, Partridge L (2008) Stress-response hormesis and aging: “That which does not kill us makes us stronger.” Cell Metabolism 7:200 – 203.

Gervasi SS, Foufopoulos J (2008) Costs of plasticity: Responses to desiccation decrease post-metamorphic immune function in a pond-breeding amphibian. Functional Ecology 22:100 – 108.

Greer AL, Berrill M, Wilson PJ (2005) Five amphibian mortality events associated with ranavirus infection in south central Ontario, Canada. Diseases of Aquatic Organisms 67:9 – 14.

Griffiths D, Kirkwood RC (1995) Seasonal variation in growth, mortality and fat stores of roach and perch in Lough Neagh, Northern Ireland. Journal of Fish Biology 47:537 – 554.

Gutzeit HO (2001) Interaction of stressors and the limits of cellular homeostasis. Biochemical and Biophysical Research 283:721 – 725.

Hamilton WJ, III (1973) Life’s Color Code. McGraw-Hill, New York, NY, U.S.A.

Hawley DM, Altizer SM (2011) Disease ecology meets ecological immunology: Understanding the links between organismal immunity and infection dynamics in natural populations. Functional Ecology 25:48 – 60.

Hayes TB, Case P, Chui S, Chung D, Haeffele C, Haston K, Lee M, Mai VP, Marjuoa Y, Parker J, Tsui M (2006) Pesticide mixtures, endocrine disruption, and amphibian declines: Are we underestimating the impact? Environmental Health Perspectives 114:40.

Heinrich B (1977) Why have some animals evolved to regulate a high body temperature? The American Naturalist 111:623 – 640.

Hirshfield MF, Tinkle DW (1974) Natural selection and evolution of reproductive effort. Proceedings of the National Academy of Sciences of the United States of America 72:2227 – 2231.

Holling CS (1959) Some characteristics of simple types of predation and parasitism. Canadian Entomologist 91:385–398.

Houlahan JE, Findlay CS, Schmidt BR, Meyer AH, Kuzmin SL (2000) Quantitative evidence for global amphibian population declines. Nature 404:752 – 755.

Huey RB, Slatkin M (1976) Costs and benefits of lizard thermoregulation. Quarterly Review of Biology 51:363 – 384.

James SM, Little EE (2003) The effects of chronic cadmium exposures on American toad (*Bufo americanus)* tadpoles. Environmental Toxicology and Chemistry 22:377 – 380.

Johnson SA, Jakob EM (1999) Leg autotomy in a spider has minimal costs in competitive ability and development. Animal Behaviour 57:957 – 965.

Jones L, Gossett DR, Banks SW, McCallum ML (2010) Antioxidant defense system in tadpoles of the American Bullfrog *(Lithobates catesbeianus)* exposed to Paraquat. Journal of Herpetology 44:222 – 228.

Jones MW, Hutchings JA (2001) The influence of male parr body size and mate competition on fertilization success and effective population size in Atlantic Salmon. Heredity 86:675 – 684.

Kapahi P, Boulton ME, Kirkwood TBL (1999) Positive correlation between mammalian life span and cellular resistance to stress. Free Radical Biology and Medicine 26:495 – 500.

Kooka K, Yamamura O, Ohkubo N, Honda S (2009) Winter lipid depletion of juvenile walleye Pollock *Theragra chalcogramma* in the Doto area, northern Japan. Journal of Fish Biology 75:186 – 202.

Lazzaro BP, Galac MR (2006) Disease pathology: Wasting energy fighting infection. Current Biology 16:R964 – R965.

Le AD, Quan B, Juzytch W, Fletcher PJ, Joharchi N, Shaham Y (1998) Reinstatement of alcohol-seeking by priming injections of alcohol and exposure to stress in rats. Psychopharmacology 135:169 – 174.

Leary CJ, Jessop TS, Garcia AM, Knapp R (2004) Steroid hormone profiles and relative body condition of calling and satellite toads: Implications for proximate regulation of behavior in anurans. Behavioral Ecology 15:313 – 320.

Lendvai AZ, Ouyang JQ, Schoenle LA, Fasanello V, Haussmann MF, Bonnier F, Moore IT (2014) Experimental food restriction reveals individual differences in corticosterone reaction norms with no oxidative costs. PLoS One 9:e110564.

Levins R (1968) Evolution in Changing Environments: Some Theoretical Explorations. Monographs in Population Biology. Princeton University Press, Princeton, NJ, U.S.A. 123 pp.

Lewis SM, Goldspink D, Phillips J, Merry B, Holehan A (1985) The effects of aging and chronic dietary restriction on whole body growth and protein turnover in the rat. Experimental Gerentology 20:253 – 260.

Li S, Tan Y (2011) Hormetic response of cholinesterase from Daphnia magna in chronic exposure to triazophos and chloryrifos. Journal of Environmental Science 23:852 – 859.

Li Y, Chen T (2005) Concentrations of additive arsenic in Beijing pig feeds and the residues in pig manure. Resources, Conservation and Recycling 45:356 – 367.

Liebig JV (1840) Organic chemistry in its application to vegetable physiology and agriculture. Readings in Ecology. Prentice Hall, New York, NY, U.S.A.

Lindstrom J (1999) Early development and fitness in birds and mammals. Trends in Ecology and Evolution 14:343 – 348.

Lochmiller RL, Deerenberg C (2000) Trade-offs in evolutionary immunology: Just what is the cost of immunity? Oikos 88:87 – 98.

Londraville RL, Macotela Y, Duff RJ, Easterling MR, Liu Q, Crespi EJ (2014) Comparative endocrinology of leptin: Assessing function in a phylogenetic context. General and Comparative Endocrinology 0:146 – 157.

MacKay WP (1985) A comparison of the energy budgets of three species of *Pogonomyrmex* harvester ants (Hymenoptera: Formicidae). Oecologica 66:484 – 494.

Manning MJ, Turner RJ (1976) Comparative Immunobiology. John Wiley & Sons, Inc. New York, NY, U.S.A.

Mattson MP (2008) Hormesis defined. Ageing Research Reviews 7:1 – 7.

Maher JM, Werner EE, Denver RJ (2013) Stress hormones mediate predator-induced phenotypic plasticity in amphibian tadpoles. Proceedings of the Royal Society B 280:20123075.

McCallum ML (2015) Vertebrate biodiversity losses point to sixth mass extinction. Biodiversity and Conservation 24:2497 – 2519.

McCallum ML (2007) Amphibian decline or extinction? Current declines dwarf background extinction rate. Journal of Herpetology 41:483 – 491.

McCallum ML (2003) Reproductive ecology and taxonomic status of Acris crepitans blanchardi with additional investigations on the Hamilton and Zuk Hypothesis. Ph.D. Dissertation. Arkansas State University, Jonesboro, U.S.A. 150 pp.

McCallum ML, Trauth SE (2007) Physiological trade-offs between immunity and reproduction in the northern cricket frog *(Acris crepitans)*. Herpetologica 63:269 – 274.

McCallum ML, Brooks C, Mason R, Trauth SE (2011) Growth, reproduction and life span in Blanchard’s Cricket Frog *(Acris blanchardi)* with notes on the growth of the Northern Cricket Frog *(Acris crepitans)*. Herpetology Notes 4:25 – 35.

McCallum ML, Matlock M, Treas J, Safi B, Sanson W, McCallum JL (2013) Endocrine disruption of sexual selection by an estrogenic herbicide in the mealworm beetle, *Tenebrio molitor*. Ecotoxicology 22:1461 – 1466.

McCallum ML, Trauth SE, McDowell C, Neal RG, Klotz TL (2006) Calling site characteristics of the Illinois Chorus Frog *(Pseudacris streckeri illinoensis)* in northeastern Arkansas. Herpetological Natural History 9:195 – 198.

McClure CD, Zhong W, Hunt VL, Chapman FM, Hill FV, Priest NK (2014) Hormesis results in trade-offs with immunity. Evolution 68:2225 – 2233.

McDiarmid RW, Altig RA (1999) Tadpoles: The Biology of Anuran Larvae. University of Chicago Press. Chicago, Illinois, U.S.A. 444 pp.

McNab BK (2002) The Physiological Ecology of Vertebrates: A View From Energetics. Comstock Publishing, Cornell University Press. Ithaca, New York, U.S.A. 587 pp.

Norris K, Evans MR (2000) Ecological immunology: Life history trade-offs and immune defense in birds. Behavioral Ecology 11:19 – 26.

Parrish JK, Edelstein-Keshet L (1999) Complexity, pattern, and evolutionary trade-offs in animal aggregation. Science 284:99 – 101.

Pastoret PP, Griebel P, Bazin H, Govaerts A (eds.). (1998) Handbook of Vertebrate Immunology. Academic Press. Cambridge, Massachusetts, U.S.A.

Perez-Coll C, Sztrum AA, Herkovits J (2008) Nickel tissue residue as a biomarker of sub-toxic exposure and susceptibility in amphibian embryos. Chemosphere 74:78 – 83.

Pianka ER (1985) Some intercontinental comparisons of desert lizards. National Geographic Research 1:490 – 504.

Pianka ER (1986) Ecology an Natural History of Desert Lizards. Analyses of the Ecological Niche and Community Structure. Princeton University Press, Princeton, NJ, U.S.A.

Pianka ER (1993) The many dimensions of a lizard’s ecological niche. Pp. 121 – 154 In: Valakos E, Bohem W, Perez V, Maragou P (eds.). Lacertids of the Mediterranean Region. A Biological Approach. Hellenic Zoological Society. Athens, Greece.

Plaistow SJ, Shirley C, Collin H, Cornell SJ, Harney ED (2015) Offspring provisioning explains clone-specific maternal age effects on life history and life span in the water flea, *Daphnia pulex*. The American Naturalist 186:376 – 389.

Pond CM (2011) Ecology of storage and allocation o fresources: Animals. In: eLS. John Wiley & Sons, Ltd. Chichester, UK.

Qiu X, Brown K, Hirschey MD, Verdin E, Chen D (2010) Calorie restriction reduces oxidative stress by SIRT3-mediated SOD2 activation. Cell Metabolism 12:662 – 667.

Quinteiro-Filho WM, Goes AVS, Pinheiro ML, Ribeiro A, Ferraz-de-Paula V, Astolfi-Ferreira, Ferreira P, Palermo-Neto J (2012) Heat stress impairs performance and induces intestinal inflammation in broiler chickens infected with *Salmonella* enteritidis. Avian Pathology 41:421 – 427.

Radak Z, Chung HY, Goto S (2005) Exercise and hormesis: Oxidative stress-related adaptation for successful aging. Biogerontology 6:71 – 75.

Ramsey JP, Reinert LK, Harper LK, Woodhams DC, Rollins-Smith LA (2010) Immune defenses against *Batrachochytrium dendrobatidis*, a fungus linked to global amphibian declines, in the South African Clawed Frog, *Xeopus laevis*. Infection and Immunity 78:3981 – 3992.

Randall JB (1981) Regeneration and autotomy exhibited by the black widow spider, *Latrodectus variolus* Walckenaer. Development Genes and Evolution 190:230 – 232.

Reed JM, Doerr PD, Walters JR (1986) Determining minimum population sizes for birds and mammals. Wildlife Society Bulletin 14:255 – 261.

Ribas L, Li M, Doddington BJ, Robet J, Seidel JA, Kroll JS, Zimmerman LB, Grassly NC, Garner TWJ, Fisher MC (2009) Expression profiling the temperature-dependent amphibian response to infection by *Batrachochytrium dendrobatidis*. PLoS One 4:e8408.

Ricklefs RE, Wikelski M (2002) The physiology/life-history nexus. Trends in Ecology and Evolution 17:462 – 468.

Schloegel LM, Picco AM, Kilpatrick AM, Davies AJ, Hyatt, Daszak P (2009) Magnitude of the US trade in aphibians and presence of *Batrachochytrium dendrobatidis* and ranavirus infection in imported North American bullfrogs *(Rana catesbeiana*). Biological Conservation 142:1420 – 1426.

Schmid-Hempel P, Schmid-Hempel P (2011) Evolutionary parasitology: The integrated study of infections, immunology, ecology and genetics. Oxford University Press. Oxford, UK. 516 pp.

Schreck CB (2010) Stress and fish reproduction: The roles of allostasis and hormesis. General and Comparative Endocrinology 165:549 – 556.

Schulenburg H, Kurtz J, Moret Y, Siva-Jothy M (2009) Introduction. Ecological Immunology. Philosophical Transactions of the Royal Society of London, Biological Sciences 364:3 – 14.

Schulz H (1888) Ueber Hefegifte. Pfluegers Arch Gesamte Physiol Menschen Tiere 42:517.

Serrano-Meneses MA, Cordoba-Aguilar, Mendez V, Layen SJ, Szekely T (2007) Sexual size dimorphism in the American Rubyspot: Male body size predicts male competition and mating success. Animal Behavior 73:987 – 997.

Sheldon BC, Verhulst S (1996) Ecological immunology: Costly parasite defenses and trade-offs in evolutionary ecology. Trends in Ecology & Evolution 11:317 – 321.

Shen K, Shen C, Lu Y, Tang X, Zhang C, Chen X, Shi J, Lin Q, Chen Y (2009) Hormesis response of marine and freshwater luminescent bacteria to metal exposure. Biological Research 42:183 – 187.

Sheridan MA (1994) Regulation of lipid metabolism in poikilothermic vertebrates. Comparative Biochemistry and Physiology Part B: Comparative Biochemistry 107:495 – 508.

Shine R (1988) The evolution of large body size in females: A critique of Darwin’s “Fecundity Advantage Model.” The American Naturalist 131:124 – 131.

Stallings CD, Coleman FC, Koenig CC, Markiewicz DA (2010) Energy allocation in juveniles of a warm-temperate reef fish. Environmental Biology of Fishes 88:389 – 398.

Stanier MW, Mount LE, Bligh J (1984) Energy Balance and Temperature Regulation. Cambridge University Press. New York, NY, U.S.A. 153 pp.

Stark JD, Banks JE, Vargas R (2004) How risky is risk assessment: The role that life history strategies play in susceptibility o fspecies to stress. Proceedings of the National Academy of Sciences of the United States of America 101:732 – 736.

Stearns SC (1989) Trade-offs in life history evolution. Functional Ecology 3:259 – 268.

Stearns SC (1992) The evolution of life histories. Oxford University Press. Oxford, UK.

Tahara EB, Cunha FM, Basso TO, Bianca BED, Gomert AK, Kowaltowski AJ (2013) Calorie restriction hysteretically primes aging Saccharomyces cerevisiae toward more effective oxidative metabolism. PLoS One 8:e56388.

Toft CA (1980) Feeding ecology of thirteen syntopic species of anurans in a seasonal tropical environment. Oecologia 45:131 – 141.

Wagner WE (1989) Fighting, assessment, and frequency alteration in Blanchard’s Cricket Frog. Behavioral Ecology and Sociobiology 25:429 – 436.

Wells KD (1977) Territoriality and male mating success in the Green Frog *(Rana clamitans)*. Ecology 58:750 – 762.

Wieser W (1991) Limitations of energy acquisition and energy use in small poikilotherms: Evolutioanry implications. Functional Ecology 5:234 – 240.

Wijesinghe MR (2012) Toxic effects of pesticides: Empirical trials provide some indication of the imminent threats to amphibians. FrogLog 20:30 – 31.

Williams GC (1957) Pleiotropy, natural selection and evolution of senescence. Evolution 11:398 – 411.

Williams GC (1966) Adaptation and Natural Selection. Princeton University Press, Princeton. U.S.A.

Young RA (1976) Fat, energy, and mammalian survival. Integrative and Comparative Biology 16:699 – 710.

Yu BP (1996) Aging and oxidative stress: modulation by dietary restriction. Free Radical Biology and Medicine 21:651 – 668.

Zahavi A (1975) Mate selection – a selection for a handicap. Journal of Theoretical Biology 53:205 – 214.

Zahavi A, Zahavi A (1999) The Handicap Principle: A Missing Piece of Darwin’s Puzzle. Oxford University Press. Oxford, U.K. 286 pp.

Zalizniak L, Nugegoda D (2006) Effect of sublethal concentrations of chlorpyrifos on three successive generations of *Daphnia carinata*. Ecotoxicology and Environmental Safety 64:207 – 214.

Zellner DA, Loaiza S, Gonzalez Z, Pita J, Morales J, Pecora D, Wolf A (2006) Food selection changes under stress. Physiology & Behavior 87:789 – 793.

Zulkifli I, Abdullah N, Azrin NM, Ho YW (2000) Growth performance and immune response of two commercial broiler strains fed diets containing *Lactobacillus* cultures and oxytetracycline under heat stress conditions. British Poultry Science 41:593 – 597.

